# Yeast Rai1p and the RAC complex (Ssz1p/Zuo1p) modulate uORF-mediated *GCN4* translational control under stress conditions

**DOI:** 10.64898/2026.03.11.711237

**Authors:** Kristína Jendruchová, Adriana Šubrtová, Leoš Shivaya Valášek

## Abstract

Translation reinitiation (REI) is one of the most important gene-specific regulatory mechanisms by which eukaryotic cells influence expression of main translons, for example during highly conserved integrated stress response (ISR). In *S. cerevisiae*, expression of the key stress response gene, *GCN4*, is controlled by an intricate interplay among four short upstream translons (uTranslons, formerly uORFs), resulting in high or low levels of REI at *GCN4* depending on the growth conditions. Under nutrient rich conditions, *GCN4* expression is repressed, but upon amino acid starvation, it is derepressed, despite of a general translational shut down. Capitalizing on our screening reporter system, we identified three new factors influencing efficiency of REI after translation of *GCN4* uTranslons: Rai1p (an RNA quality control and processing factor), and Ssz1p and Zuo1p (members of the Ribosome Associated Complex [RAC]). Importantly, we showed that depletion of these factors deregulated derepression of Gcn4p synthesis under starvation. Furthermore, we found that similar to RAC, Rai1p associates with 40S subunits and actively translating ribosomes. We also explored interactomes of these proteins. Collectively, we present three previously unknown factors that co-regulate stress response to amino acid starvation in the budding yeast by unique mechanisms.

## INTRODUCTION

Translation is a tightly regulated step of gene expression. It is influenced by a network of *cis*- and *trans*-acting factors that all act in concert to promote the action of the main actor in this process – the ribosome. This molecular machine is comprised of the small 40S subunit and large 60S subunit that join together to form the elongation-competent 80S ribosome. Translation initiation naturally begins the whole translational cycle, for which the translation initiation site (TIS; in the most cases AUG) of mRNA must be recognized by the anticodon of methionyl initiator tRNA (Met-tRNA_i_^Met^). Translation of the most eukaryotic mRNAs depends on the so-called scanning mechanism, where the 40S subunit is first loaded with Met-tRNA_i_^Met^ in a ternary complex (eIF2-TC) with GTP-bound eukaryotic Initiation Factor (eIF) 2 in a reaction promoted by eIF1, eIF1A, eIF5 and the multisubunit complex eIF3. This forms the 43S pre-initiation complex (PIC), which in turn is recruited to the 5’ end of the mRNA by the eIF4F complex, forming the 48S PIC. The 5’ UTR of the mRNA is then inspected base-by-base for complementarity to the anticodon of Met-tRNA_i_^Met^ as the individual mRNA codons enter the 40S decoding site. Subsequent TIS recognition triggers GTP hydrolysis on eIF2, followed by the release of eIF2-GDP and eIF5 from the 40S subunit. The whole process is captured by the 60S joining step followed by GTP hydrolysis on eIF5B. Upon ejection of eIF5B-GDP coupled with eIF1A, the 80S initiation complex carries on into elongation (reviewed in^1,2,3^). Completion of a synthesis of a given protein is marked by translation termination, followed by a 2-step ribosome recycling phase to allow a new round of translation to begin.

However, in specific cases termination is followed by only a partial 60S recycling (i.e. only the first recycling step materializes), which enables another 40S-mediated initiation event to occur downstream of the just translated translon on the same mRNA molecule. (Translon [short for translated region] refers to a recently proposed alternative to the Open Reading Frame [ORF] and Coding DNA Sequence [CDS], denoting any region that is genuinely decoded by the ribosome^4^. This process is called translation reinitiation (REI) (reviewed in^3,5^). REI is often utilized by the cell to regulate the rate of translation of the main translon through translation of short upstream translons (uTranslons, formerly uORFs) located in the 5’ UTR region of an mRNA. Translation of uTranslons is by definition inhibitory towards translation of the main translon^3^. Importance of this type of translational regulation is illustrated by the fact that around ∼49% of human^6^ and ∼13% of yeast^7^ transcripts contain uTranslons.

Mechanistically, REI can be achieved via the following ways: (i) by the partial post-termination complex (post-TC) recycling mentioned above, where the 60S subunit is removed by the action of ABCE1/Rli1p, which is followed by either spontaneous or Tma22p/DENR–Tma20p/MCTS1-mediated release of deacylated tRNA from the ribosomal P-site^8,9^; or (ii) by blocking the whole recycling process, in which case REI may occur with the complete post-TC; i.e. with both subunit still bound to each other, in a manner that may or may not require release of the deacylated tRNA^3,5,10^.

The rate of REI is regulated by an array of *cis*- and *trans*-acting factors, including uTranslons and other specialized mRNA features, as well as various protein effectors. For instance, uTranslon length was demonstrated to inversely correlate with the REI efficiency^11–13^, which was later attributed to post-initiation retention of some eIFs, like eIF3 and eIF4G, by 80S ribosomes translating shorter uTranslons^3,14–17^. It was demonstrated that the efficiency of this unexpected retention, and thus the REI efficiency, gradually decreases with an increasing length of translated uTranslon. On the other hand, REI efficiency increases with the increasing distance between the uTranslon stop codon and the main translon TIS^18,19^. This is due to the increasing probability of the eIF2-TC reacquisition by the post-termination 40S over the longer distance that it has to traverse to become fully competent of recognizing the next TIS^19–21^.

Hypothetically, ribosome recycling factors might also contribute to changes in REI efficiency. After the splitting of both ribosomal subunits by Rli1p/ABCE1 and removal of deacylated tRNA from the P-site by Tma22p/DENR–Tma20p/MCTS1 and/or by Tma64p/eIF2D, the latter factors also separate the 40S subunit from the mRNA^10,16,22,23^. Indeed, their absence induces genome-wide accumulation of unrecycled 40S subunits at stop codons in both uTranslons and main translons, which is biased by certain penultimate codons over the others^23,24^. It was therefore hypothesized that disassociation of only some tRNAs, i.e. those corresponding to these penultimate codons, is strongly dependent on the action of the 40S-recycling factors^9,23,24^. Accordingly, only the “Tma/DENR-dependent” codons showed significantly greater accumulation of unrecycled 40S subunits. It is important to note that even the least Tma-dependent codons in yeast showed at least ∼4-fold greater 40S accumulation levels in the deletion strains compared to WT^24^. Importantly, however, we have recently demonstrated that even though these codon preferences do apply to bulk 40S recycling, they do not translate into an equivalent change in the rate of REI in the absence of the Tma factors. In other words, this Tma-dependency does not impose the same rules for the efficiency of REI^25^.

Arguably, the most prominent example of translational control via regulatory uTranslons is the so-called “delayed REI”. This mechanism governs translation of Gcn4p in *S. cerevisiae*, as the most thoroughly studied case. Gcn4p is a transcription factor that regulates the yeast’s response to amino acid starvation through activation of genes falling under the General Amino Acid Control (GAAC) pathway (reviewed in ^26,27^). The *GCN4* 5’UTR contains four short uTranslons (originally called uORF1 – 4; therefore we are not changing their designation), and translation of its mRNA is very sensitive to eIF2-TC levels, which vary depending on different nutrient conditions. In particular, uORF1 is efficiently translated under any conditions, and since it is generally permissive for REI, it prevents the 2^nd^ recycling step to allow the 40S post-TC to start traversing downstream. In non-stressed cells (with high eIF2-TC levels), most traversing ribosomes reacquire the eIF2-TC before reaching the REI-non-permissive uORFs 3 or 4, therefore ribosomes terminating at these two uTranslons undergo full recycling and never reach the main *GCN4* translon. In contrast, under starvation conditions, the Gcn2p kinase senses low amino acid levels and phosphorylates the alpha subunit of eIF2. This suspends the formation of new eIF2-TCs and, as consequence, downregulates global translation rates. As a result, ribosomes traversing downstream of uORF1 are largely unable to reacquire the new eIF2-TC before reaching uORFs 3 and 4, effectively bypassing their TISs. Critically, a significant proportion of these ribosomes reacquires the eIF2-TC while on their way from uORF4 to the *GCN4* TIS, which triggers Gcn4p expression^19,28,29^. A relatively very similar mechanism operates in mammals via the master regulator of mammalian integrated stress response – ATF4^30^, although certain differences involving various *cis*-acting elements have been recently reported by us and others^31–36^.

Mechanistically, it has been determined that uORF1 and functionally similar uORF2^19,37^ are highly REI permissive due to the so-called REI-promoting elements (RPEs) forming the 5’ enhancer, which functional interacts the eIF3a/TIF32 subunit of eIF3^20,21,37,38^ (Figure 1A). REI is also potentiated by sequences immediately following uORF1 and by penultimate codon identity of uORFs 1 and 2^19,39^. In contrast, the REI-non-permissiveness of uORF3 and uORF4 is given by both their codons identity and their 3’ nucleotide context (Figure 1). Note that REI efficiency of uORF4 is about 3-5% of that of uORF1^20,37^.

**Figure 1.**
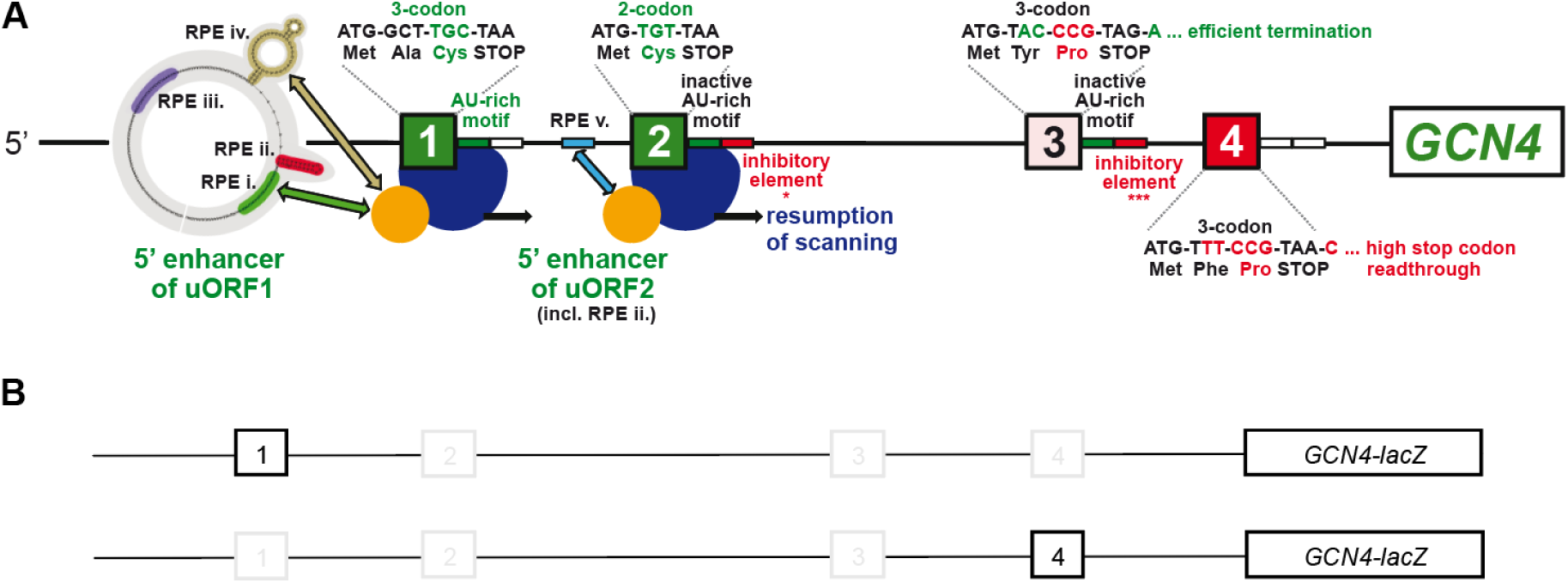
Summary of all *cis*-determinants that either promote or inhibit reinitiation on *GCN4* after translation of its four short uTranslons (uORFs). **A.** Green color-coding indicates stimulatory effects of the corresponding *cis*-factors on efficiency of REI, whereas red color-coding indicates inhibitory effects (with the exception of RPE ii. of uORF1, which is also stimulatory); the number of asterisks below the inhibitory elements of the uORF2 and uORF3 3′ sequences illustrates the degree of their inhibition. Adapted with permission from^19^. **B.** Schematic of the uORF1-only (top) and uORF4-only (bottom) constructs used in our reporter setup. uORFs with mutated TIS for each construct are greyed out.

Building upon this knowledge, we used a well-established *GCN4*-based reporter system^25^ as a genetic tool to systematically search for new factors influencing efficiency of REI and ribosome recycling after translation of short uTranslons. In particular, we examined the REI rate after REI-permissive uORF1 and REI-non-permissive uORF4 in yeast strains deleted for individual candidate factors compared to wild-type. This robust screening identified: i.) the decapping endonuclease Rai1p, and ii.) the Ssz1p/Zuo1p heterodimer, which forms the Ribosome Associated Complex (RAC). Besides decapping, Rai1 (DXO in mammals) has a role in mRNA surveillance and ribosome biogenesis, among others^40–42^. The RAC complex is known to associate with ribosomes^43–46^ and controls nascent peptide folding^43^, -1 frameshift^45^ and stop codon readthrough (SC-RT)^47^. Our genetic and biochemical analysis revealed that these factors modulate both *GCN4* translational control and general translation.

## RESULTS

### Rai1p and the RAC complex influence REI after translation of uORF1 and uORF4

To screen our pre-selected candidates (see below), we used reporter plasmids containing ∼600 nt long 5’ UTR of the *GCN4* gene followed by ∼150 nt of its translon fused in-frame with the *LacZ* reporter gene. To create a single uORF1 or uORF4 reporters, the AUG start sites of other three uTranslons where eliminated by point mutations in given constructs. In particular, the “uORF1” construct^48^ bears only REI-permissive uORF1, while the “uORF4” construct^28^ carries just REI-non-permissive uORF4 (Fig. 1B). This reporter system has been successfully used in the past to dissect the specifics of REI and ribosome recycling on *GCN4* mRNA, and thus to decipher its translational control^19–21,25,26,28,37,39,48^. Inferring from this wealth of knowledge, we applied the following logic in our study. It was shown that REI defects manifest as lower REI rates after translation of uORF1, resulting in lower “uORF1” reporter activity^20,21^. Thus, mutations or gene deletion impairing REI should lower the “uORF1” reporter activity compared to WT, warranting further analysis. On the other hand, ribosome recycling defects generally manifest as higher REI activity after translation of uORF4 (using the “uORF4” reporter compared to WT), since lower rates of ribosome recycling allow more 40S post-termination ribosomes to remain mRNA-bound even after translation of REI-non-permissive uORF4^25^. We performed our screening experiments in the BY4741 genetic background, utilizing strains obtained from the Yeast Deletion Collection^49^.

The rationale for selecting our candidate genes was as follows. We selected genes from different classes of translation-related factors that had already been implicated in promoting either REI or recycling based on *in vitro* and/or indirect evidence. In parallel, we picked factors associated with different translation processes whose connection to ribosome recycling or REI could be predicted based on their known or proposed cellular functions. Among others, the selected gene groups included initiation factors, termination and related factors, RNA helicases, ribosome and/or translation-associated ATPases or GTPase and related proteins, factors associated with *GCN4* regulation, and finally, numerous as-yet-uncharacterized genes encoding proteins that co-purified with yeast ribosomes in proteomic screens, with our particular interest put on 12 proteins designated as translation-machinery-associated (TMA)^50^. (The list of candidates that passed the initial screening process is provided in Table S1.)

Our thorough screening identified three top candidates that matched our most stringent criteria: Ssz1p and Zuo1p (together forming the RAC complex), and Rai1p for validation and further analysis. There was no significant difference in *GCN4-LacZ* expression after uORF1 translation in *ssz1Δ* and *zuo1Δ* deletions strains compared to WT (Fig. 2A), however, we found a significant >1.5-fold increase in the uORF4 reporter activity in both RAC deletion strains (Fig. 2B), suggesting a recycling defect. Even though Rai1p was not known to function in cytoplasmic translation, our reporter analysis showed a ∼50% reduction in the uORF1 reporter activity in the *rai1Δ* deletion strain (Fig. 2C). It also displayed an apparent, although not statistically significant in our experimental set-up, drop in the uORF4 activity (Fig. 2D); i.e., we recorded virtually the opposite phenotype to that seen with the RAC proteins with this particular uTranslon. Therefore, based on our logic, Rai1p most probably promotes REI. Given the REI-non-permissive character of uORF4, the observed statistical insignificance is not surprising because the Rai1p stimulatory effect must be small at this uORF.

**Figure 2.**
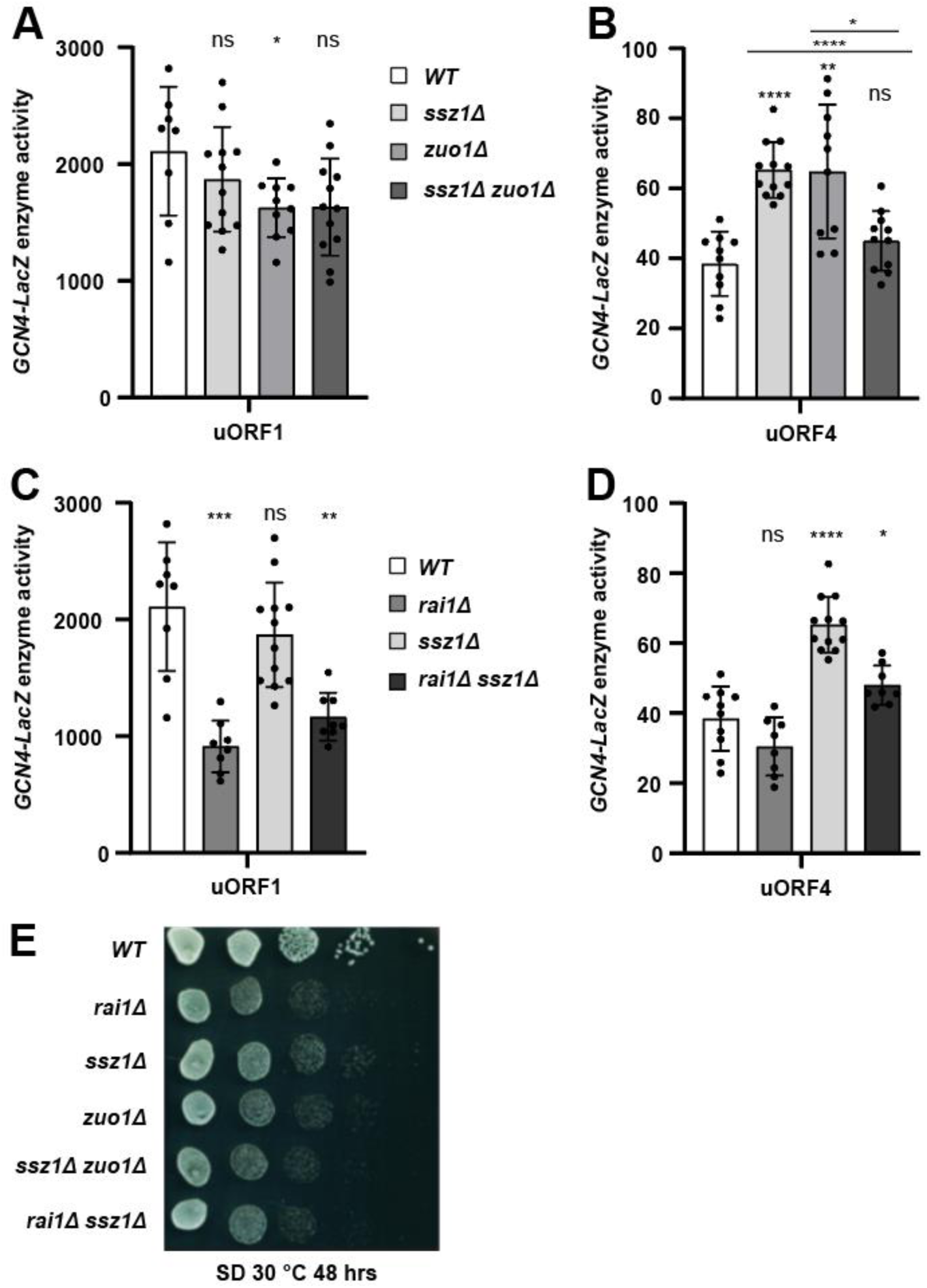
Impact of *rai1*, *ssz1,* and *zuo1* single and double deletion on *GCN4-lacZ* reporter expression and cell growth. **A**. WT, *ssz1Δ*, *zuo1Δ* and *ssz1Δ zuo1Δ* yeast strains were transformed with a plasmid containing uORF1 followed by the *GCN4-LacZ* fusion. Reporter activity was assayed in at least 5 independent transformants and normalized as described in Methods and Materials. Data are presented as normalized mean values ± SD with points denoting each individual reporter value. Statistical analysis was performed using two-tailed unpaired Welch’s t test. \**p* < 0.05, \*\**p* < 0.01, \*\*\**p* < 0.001, \*\*\*\**p* < 0.0001, ns - not significant. For source data of the measurements, please see Supplementary Data 1. **B.** Same as in (A), but the strains were transformed with the uORF4 reporter plasmid. **C-D.** Same as in (A-B), only WT, *rai1Δ*, *ssz1Δ* and *rai1Δ ssz1Δ* yeast strains were examined. **E.** The indicated yeast strains were grown overnight, diluted to an OD_600_ = 0.5, then serially diluted (10x) and spotted onto SD plates. The plates were incubated at 30 °C for 48 hours

Since Zuo1p and Ssz1p form a heterodimer, we created and tested a double deletion strain. Interestingly, loss of both RAC subunits led to the loss of the increased uORF4 phenotype with no impact on uORF1 (Fig. 2A and B). Informed by a yeast proteome survey reporting a physical interaction between Rai1p and Ssz1p^51^, we also created and tested the *rai1Δ ssz1Δ* double deletion strain. This strain exhibits a significant, nearly as robust decrease (almost 2-fold) in the uORF1 reporter expression as the single *rai1Δ* strain (Fig. 2C). Strikingly, *rai1Δ ssz1Δ* displays only modest (but still significant) increase in the uORF4 reporter expression, which is practically an average activity of the *rai1Δ* and *ssz1Δ* individual activities combined (Fig. 2D). These phenotypes suggest: i) a dominant uORF1-associated REI phenotype of *rai1Δ versus ssz1Δ*; and ii) a compensatory uORF4-associated REI phenotype of both single deletions. We failed to create a *rai1Δ zuo1Δ* strain despite many tries.

Collectively, loss of Rai1p hampers efficiency of REI after translation of uORF1 (and to some degree also after uORF4) with no effect exerted by the RAC complex at this uTranslon, whereas the loss of RAC complex impairs ribosome recycling at uORF4 (thus allowing increased REI past this uTranslon), which can be nearly fully rescued by a concomitant loss of Rai1p. These findings might point to a possible physical interaction between Raip1 and the RAC components (see below).

### Rai1p and the RAC complex modulate *GCN4* translational control

Next, we analyzed the growth phenotype of WT and all deletions and their combinations using the yeast spot assay (Figure 2E). All single deletion strains exhibited a slow growth phenotype (Slg^-^) at 30 °C on SD media, underscoring the importance of these factors for cell growth and fitness. The phenotype was more severe in the case of *rai1Δ* and *zuo1Δ* compared to *ssz1Δ*, but no additive or compensatory phenotypes were observed (Fig. 2E).

The key experiment was to test whether deletions of *rai1Δ*, *zuo1Δ*, and *ssz1Δ* perturb general translational control of *GCN4* under amino acid stress conditions induced by 3-aminotriazole (3-AT), because only if they do, their REI-associated role in the cell is physiologically significant. 3-AT functions as a competitive inhibitor to imidazole glycerol-phosphate dehydratase (encoded by the *HIS3* gene), which acts at the last step of the histidine biosynthesis pathway^52,53^. Lack of histidine leads to accumulation of uncharged histidyl-tRNAs, which activates the Gcn2p kinase that phosphorylates eIF2α, which in turn induces Gcn4p translation^29,54^. Since the BY4741 genetic background bears *his3Δ*^49^, a functional *HIS3* gene on an empty shuttling plasmid needed to be introduced to all tested strains along with our well-established reporter plasmid (B196). It contains only uORF1 and uORF4 in the *GCN4* 5’ UTR and was shown to faithfully recapitulate the complex *GCN4* translational control^28^.

The β-galactosidase reporter assays using this specific reporter plasmid expressed in WT and all deletion strains incubated with or without 10 mM 3-AT revealed the following effects (Fig. 3A): i) no significant changes were seen under normal conditions in any deletion strain; ii) all deletions or their combinations robustly reduced the *GCN4* inducibility under starvation compared to WT (the weakest but still significant defect was seen in *ssz1Δ*); iii) the compensatory uORF4-associated REI phenotype of *rai1Δ ssz1Δ* did not translate into the overall GCN4 control (highly likely due to the dominant uORF1-associated REI phenotype of *rai1Δ versus ssz1Δ*). The practically identical ratio of 3-AT-treated/untreated reporter activities in each strain can be seen in Figure 3B.

**Figure 3.**
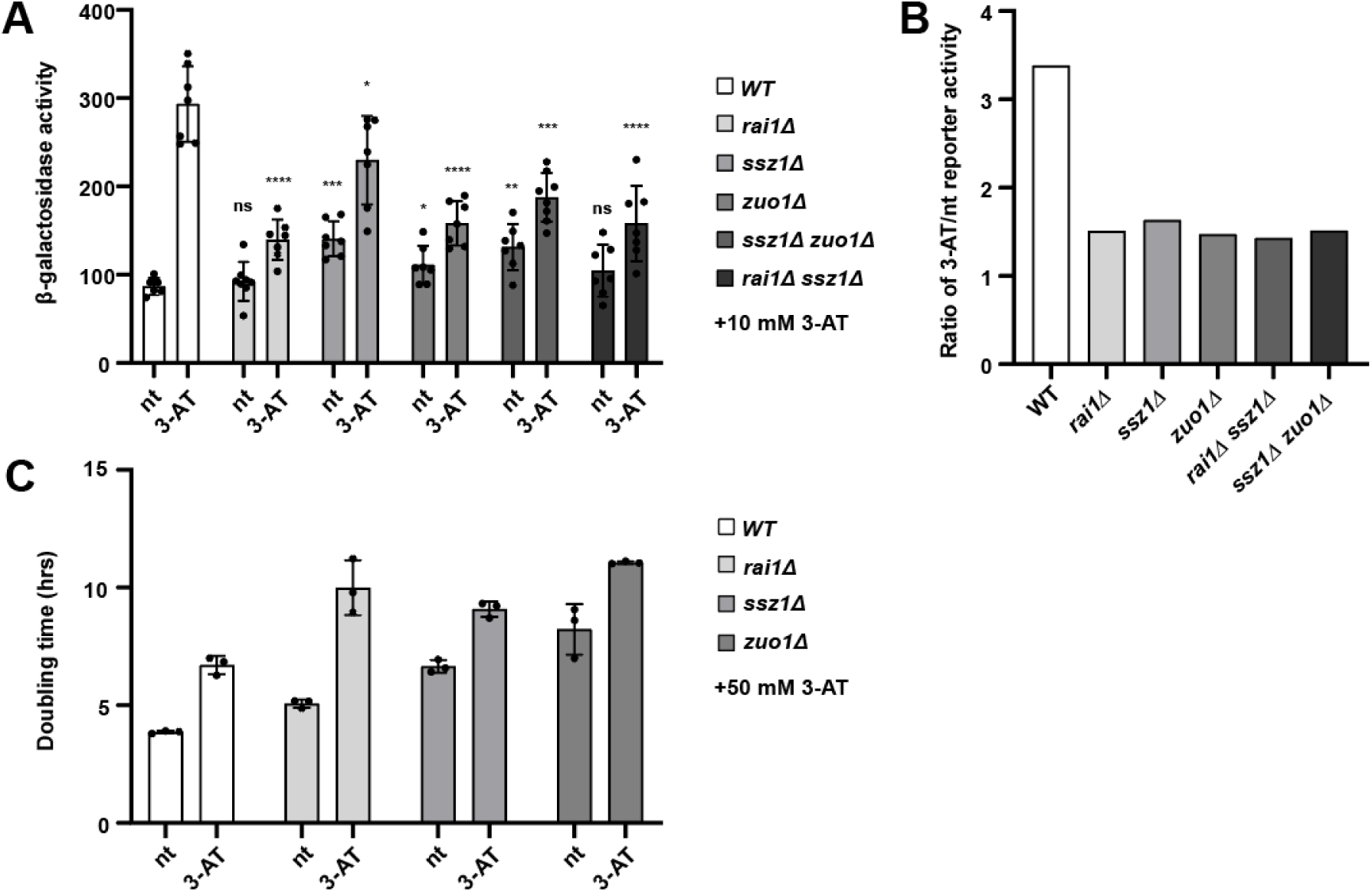
Rai1p and the RAC complex modulate *GCN4* translational control. **A**. The indicated yeast strains were co-transformed with a vector containing the *HIS3* gene and another one containing the uORF1+uORF4 reporter, and the resulting transformants were grown in parallel with or without 10mM 3-AT for 4 hours. Reporter activity was assayed in at least 5 independent transformants. Data are presented as normalized mean values ± SD with points denoting each individual reporter value. Statistical analysis was performed between WT versus deletion strains for untreated and treated values, respectively, using two-tailed unpaired Welch’s t test. \**p* < 0.05, \*\**p* < 0.01, \*\*\**p* < 0.001, \*\*\*\**p* < 0.0001, ns not significant. For source data of the measurements, please see Supplementary Data 1.**B.** Ratio of reporter activities from (A) between 3-AT-treated and untreated cultures, indicating the rate of the Gcn4p induction. **C.** Doubling time of WT, *rai1Δ*, *ssz1Δ* and *zuo1Δ* strains transformed with a plasmid containing *HIS3* grown in minimal SD media. Values are averages of 3 biological replicates with individual values indicated by data points.

Since all tested deletion strains reduced the *GCN4* inducibility by at least ∼50% (Fig. 3B), we next examined whether the low inducibility translates also into a physiologically relevant growth defect. To do that, WT and individual deletion strains transformed the *HIS3* empty plasmid were assayed for growth rates in the liquid minimal (SD) media supplemented with leucine, methionine and uracil, either supplemented with 50 mM 3-AT or not. Their doubling time (dt) was calculated, and we compared the fold-change between WT and deletion strains for untreated and 3-AT-treated cultures (Fig. 3C and D). Under normal conditions, the dt of *rai1* deletion was least affected compared to wt, followed by *ssz1* and *zuo1* deletions, while under stress conditions, the dt of all there deletions was comparable (∼1.5-fold higher than wt). Thus, the markedly reduced *GCN4* inducibility resulted in a comparably severe slow growth of all mutants, despite the fact that their starting position (dt under nt) differed. Collectively, our findings illustrate the importance of all three factors in the physiological well-being of the yeast cells, which can be at least partially attributed to the proper functioning of the delayed REI mechanism of the *GCN4* regulation.

### Rai1p associates with 40S subunits and actively translating 80S ribosomes

To assess the impact of single or double deletions of *RAI1*, *SSZ1* and *ZUO1* on general translation rates, we performed polysome profiling using cycloheximide, which freezes elongating ribosomes (Figure 4). As shown in Fig. 4B, *rai1Δ* exhibited the so-called ‘halfmers’ that can be seen as trailing shoulders of 80S monosomes and polysomes (marked with an arrow) and suggests either a 60S biogenesis defect or 60S–48S subunit joining defect. At the same time, *rai1Δ,* exhibited a mild polysome runoff (indicated by reduction in polysomal fractions) (Fig. 4B). A more pronounced runoff was seen with *ssz1Δ* and *zuo1Δ* single and *ssz1Δ zuo1Δ* and *rai1Δ ssz1Δ* double deletions, indicating a clear reduction of initiation rates (Fig. 4C - F). Interestingly, as in the case of the “uORF4 phenotype” of *ssz1Δ* that was suppressed *rai1Δ* (Fig. 2D), here, combining *rai1Δ* and *ssz1Δ* suppressed the halfmer phenotype of *rai1Δ*.

**Figure 4.**
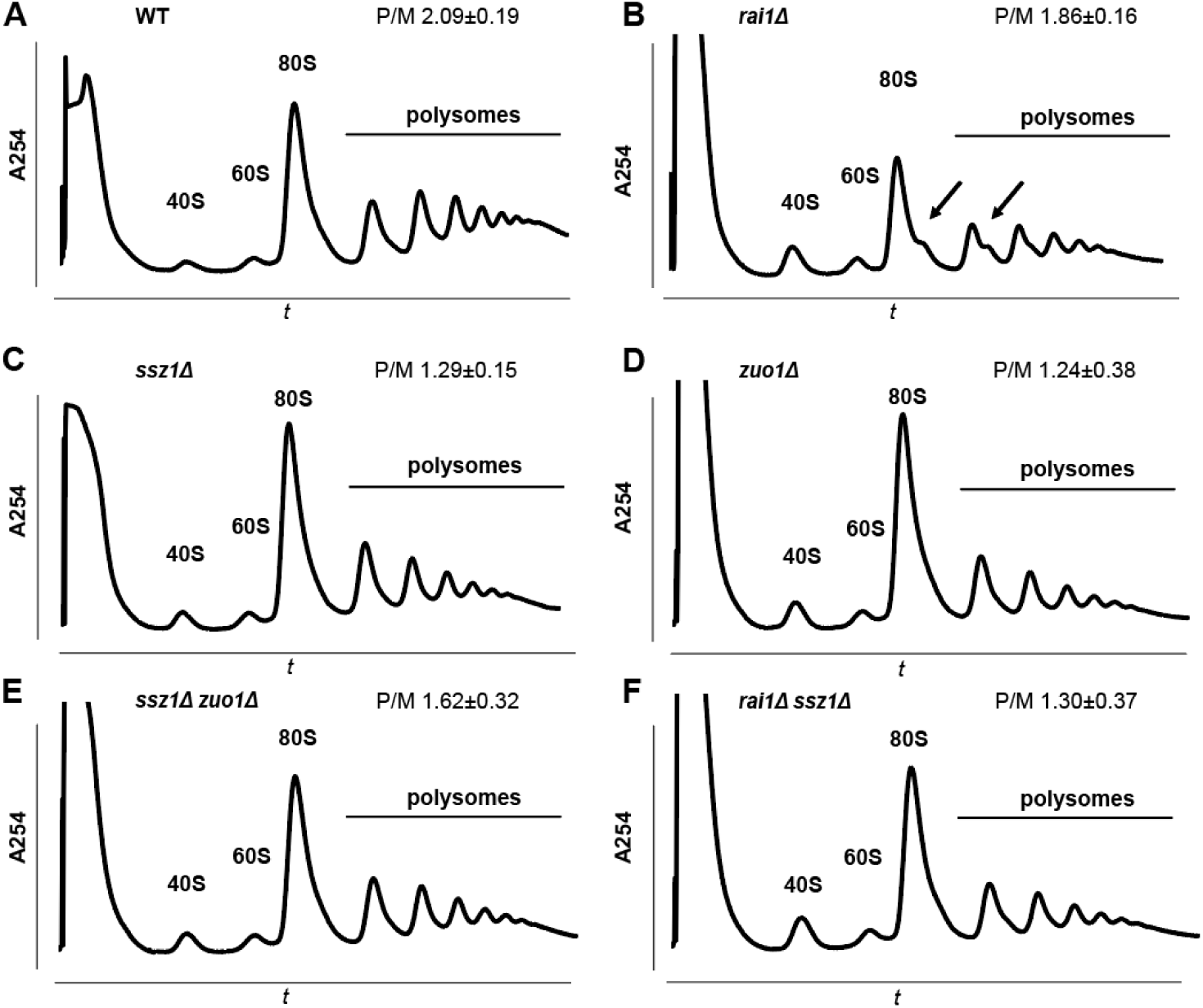
Polysome profiles of wild type and indicated deletion strains. **A –F.** Yeast strains were grown to an OD_600_ ∼ 0.8 in YPD media, treated with cycloheximide and WCEs were prepared. WCE aliquots containing 8U of total RNA were layered onto 5-50% sucrose gradients and separated by ultracentrifugation at 260,000g for 2.5 h. Gradients were scanned at 254 nm to visualize ribosomal species. Positions of 40S, 60S, 80S and polysomal species, as well as halfmers, are indicated, and P/M ratios are displayed in the upper right corner.

Next, we wished to examine whether our three candidates can reciprocally suppress Slg^-^ phenotypes associated with their deletions when in high copy. Therefore, we created overexpression plasmids carrying the *RAI1*, *ZUO1* and *SSZ1* genes fused C-terminally with a 6x Gly linker followed by 3x FLAG tag. These protein fusions were flanked by 500 nt of the natural 5’ UTR and 3’UTR contexts of their corresponding genes. Growth complementation spot assays confirmed that the FLAG-tagged fusion proteins complemented growth defects in the corresponding deletion strains (Fig. S1A – C). However, we saw no growth complementation of *rai1Δ* with either *SSZ1-FLAG* or *ZUO1-FLAG* (Fig. S1B); of *ssz1Δ* with either *RAI1-FLAG* or *ZUO1-FLAG* (Fig. S1B); and of *zuo1Δ* with either *SSZ1-FLAG1* or *RAI1-FLAG* (Fig. S1C).

Given the observed REI/recycling phenotypes, we were curious to test whether these three proteins associate with small ribosomal subunits, as could be expected from our results and some previous studies; the Ssz1p:Zuo1p heterodimer has been shown to occur on ribosomes^44–46,55^. However, nothing is known about cytoplasmic Rai1p and ribosomes. Thus, we grew *rai1Δ, ssz1Δ* and *zuo1Δ* strains transformed with the corresponding *RAI1/SSZ1/ZUO1-FLAG* low copy plasmids, fixed the cells with 1% formaldehyde as a crosslinking agent, generated and separated WCEs by high velocity centrifugation, collected fractions from sucrose gradients and performed western blotting to determine their presence in each fraction (Figure 5). The eIF3a/Tif32p subunit of eIF3, known to associate with initiating, elongating and terminating ribosomes^14,17,56–59^, was used as a control. While Rai1-FLAG was detected mainly in the top (ribosome-free) fractions, a sizeable proportion co-sedimented in the 40S-containing fractions, as well as in monosomal and even polysomal fractions, containing actively translating ribosomes (Fig. 5A). The Ssz1-FLAG and Zuo1-FLAG showed a very similar co-sedimentation pattern to Rai1p (Fig. 5B and C), consistent with the published studies^44–46,55^.

**Figure 5.**
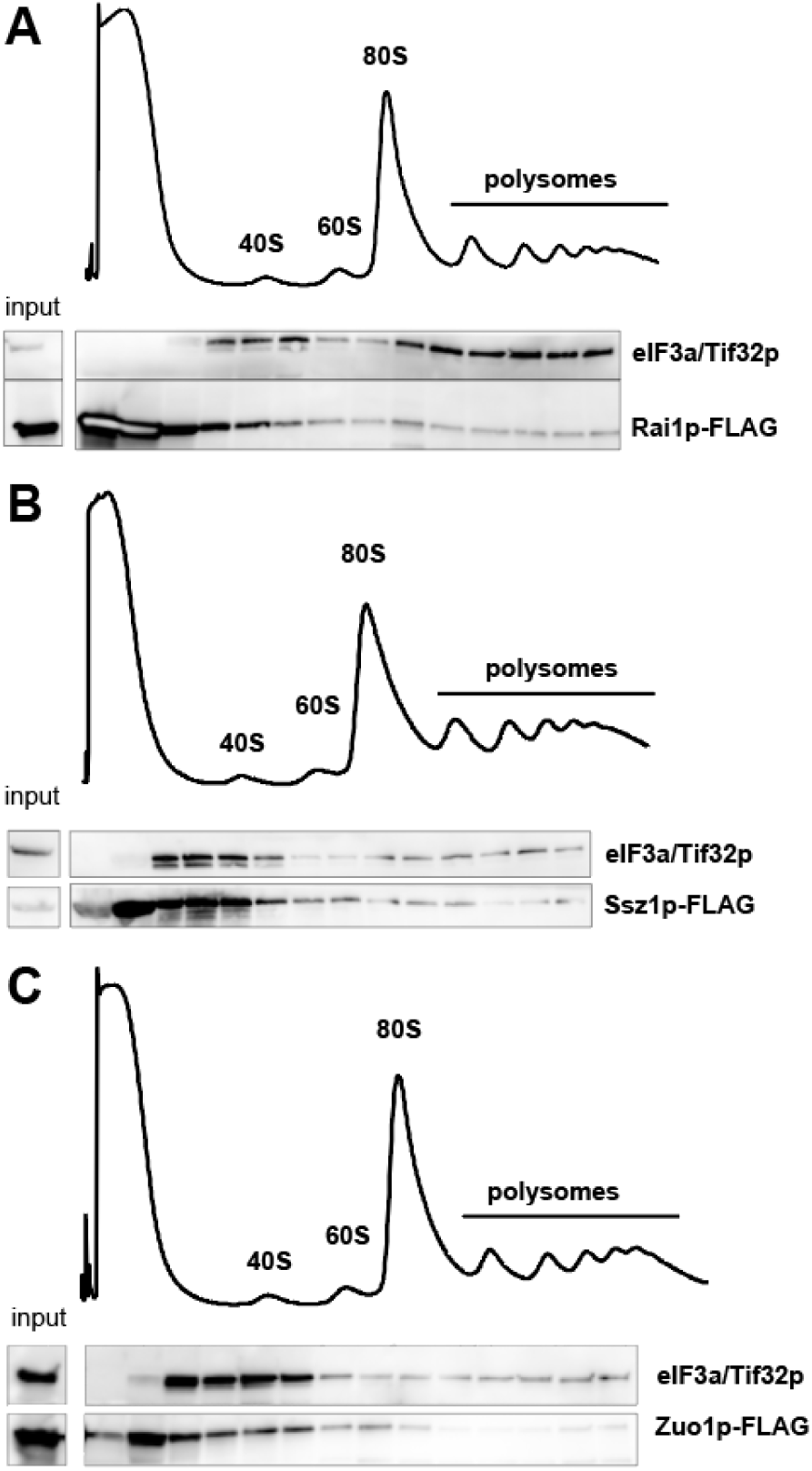
Rai1p associates with 40S subunits and actively translating 80S ribosomes. **A.** The *rai1Δ* strain transformed with YEplac181 carrying a *RAI1-FLAG* fusion was grown in SD media to an OD_600_ ∼ 0.9-1.0 and crosslinked with 1% formaldehyde. 20 U of A_260_ were layered onto 5-50% sucrose gradients and separated by ultracentrifugation at 39,000rmp for 2.5 h. Gradients were scanned at 254 nm to visualize ribosomal species, fractions were collected, precipitated with 96% EtOH, and subjected to western blot analysis. See Methods for details. **B.** Same as A, except that *ssz1Δ* transformed with YCplac111 carrying a *SSZ1-FLAG* fusion was examined. **C.** Same as A, except that *zuo1Δ* transformed with YCplac111 carrying a *ZUO1-FLAG* fusion was examined.

### Rai1p and Ssz1p interactomes

To explore an interactome of Rai1p as a newly characterized translation factor, we performed co-immunoprecipitation experiments (co-IP) of Rai1-FLAG-associated complexes. As a reference co-IP sample, we used Ssz1-FLAG-associated complexes, and as negative control, we used the *rai1Δ* or *ssz1Δ* deletion strains. The yeast cultures transformed with YEplac181 plasmids carrying FLAG-tagged fusions were grown to an OD_600_ ∼ 0.8 in minimal SD media, cells were lysed and immunoprecipitated complexes were then sent for Liquid Chromatography/Mass Spectrometry (LC/MS) analysis (Fig. 6A - B). Due to high abundance of ribosomal proteins in cells in general, which can potentially obfuscate the MS analysis^60^, we performed co-IP experiments under more stringent salt conditions. This way we could distinguish background contaminants from true interactors (see the Methods section for more details). Validating our co-IP, Rai1p pulled down Rat1p, its canonical nuclear interaction partner^42^. Among the most intriguing Rai1p-specific hits was Dbp5p, which is a DEAD-box helicase that has been implicated in RNA metabolism, translation termination and RNA quality control (reviewed in^61^). In addition, ribosomal proteins were expectedly detected, as well as several proteins from amino acid biosynthetic pathways, such as Arg1p, His4p, Hom2p and Cpa2p, which was not expected (Fig. 6A). Validating our approach further, Ssz1-FLAG brought down its RAC partner – Zuo1p – as one of the top hits (Fig. 6B). Furthermore, expected enrichment of ribosomal proteins was seen including the 40S head interacting hub – Asc1p, plus other ribosome or ribosome biogenesis-associated factors (Utp11/20p, Bcp1p, Spb1p), and some proteins involved in amino-acid synthetic pathways (Asn1/2p, Lys1p, Aat1p) (Fig. 6B). Interestingly, both co-IP experiments yielded Snz1p, a pyridoxine biosynthesis protein whose gene belongs among Gcn4p transcription factor target genes^62^ (Fig. 6A - B). Since we saw no Rai1p in the Ssz1p-FLAG co-IP and *vice versa*, we tested their prospective interaction using an *in vitro* protein-protein binding assay. Indeed, we observed a binding between GST-Ssz1p and radiolabelled Rai1p, which was weaker than the binding with its well-known interaction partner Zuo1p but still obvious (Fig. 6C). Despite the fact that none of the identified hits clearly pointed to a specific role for the studied proteins in the REI mechanism and/or ribosomal recycling, overall, our interactome data further support Rai1p’s newly described role in translation as a ribosome associated factor.

**Figure 6.**
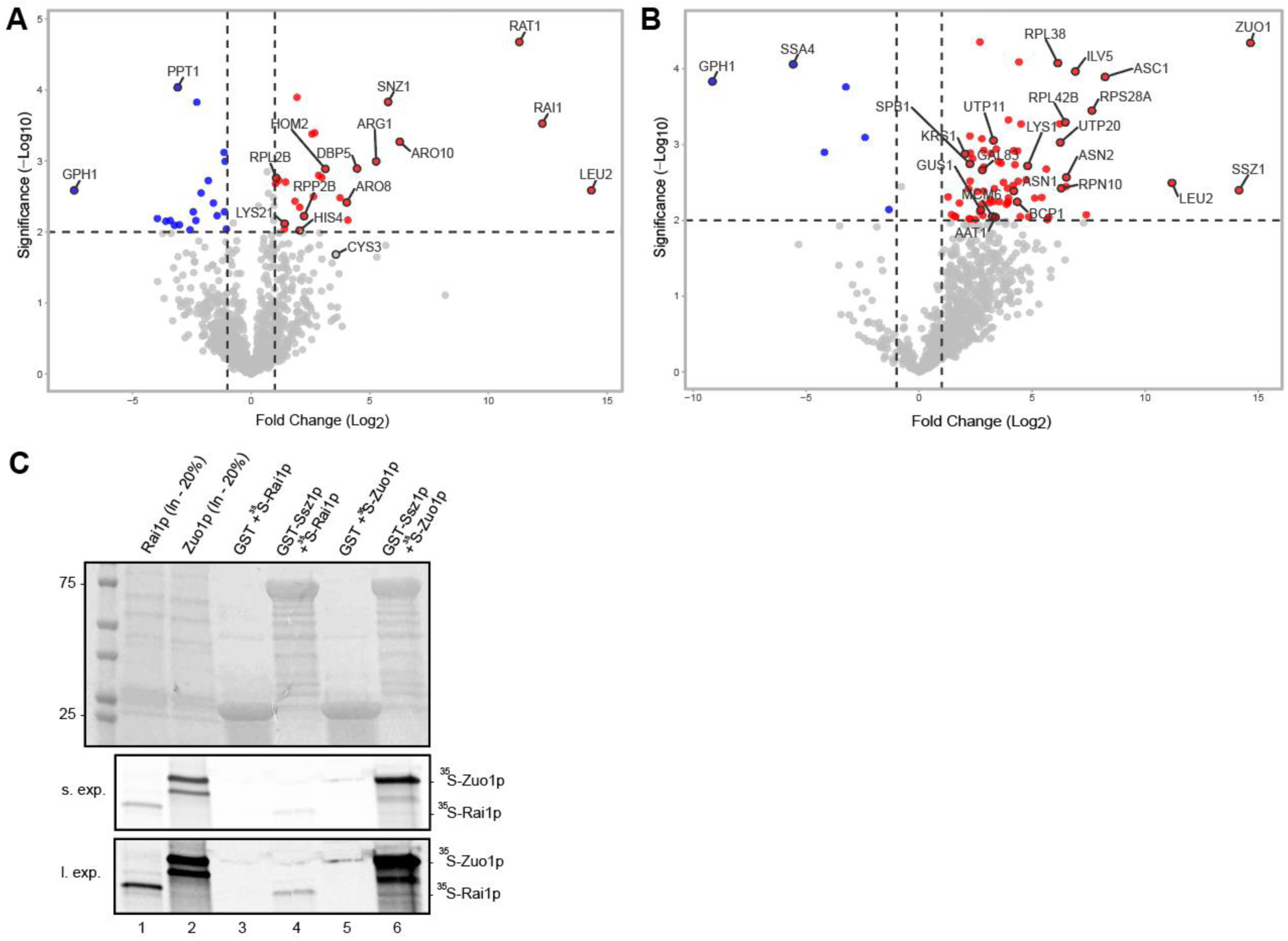
Rai1p and Ssz1p interactomes. **A - B.** Volcano plots of LC/MS analysis of complexes pulled down by (A) *RAI1-FLAG* expressed in *rai1Δ* and (B) *SSZ1-FLAG* in *rai1Δ*. As negative controls we used deletion strains. Log_2_FC threshold was set as - 1/+1 (corresponding to 2-fold change) and -Log p value threshold was set as 2 (corresponding to *p* = 0.01). **C.** Ssz1p fused to GST (lanes 4 and 6) or GST alone (lanes 3 and 5) was expressed in *E. coli*, immobilized on glutathione-Sepharose beads, and incubated with the ^35^S-labeled Rai1p (lanes 3 – 4) and Zuo1p (lanes 5 – 6) at 4 °C for 2 h. The beads were washed three times with phosphate-buffered saline, and bound proteins were separated by SDS-PAGE. Gel were first stained with GelCode Blue Stain Reagent (top panel) followed by autoradiography (bottom panels are the same but differ in exposition time). Lanes 1 and 2 show 20% of the input (In) amounts of ^35^S-labeled Rai1p and Zuo1p, respectively.

## DISCUSSION

In our long-standing effort to identify new *cis-* or *trans-*acting factors contributing to REI, we employed a *GCN4*-based reporter system and identified three new factors modulating efficiency of REI at short uTranslons – Rai1p, Ssz1p and Zuo1p. The Ssz1p/Zuo1p heterodimer forming the RAC complex has already been implicated in translational control – specifically, it influences co-translational folding^63–66^, -1 frameshifting^67^ and SC-RT^47^. The mammalian RAC complex (Hsp70L1/Ssz1 and MPP11/Zuo1), shown to complement the yeast RAC deletion, similarly modulates SC-RT^68^. We propose that the higher uORF4-only reporter activity that we observed with the *ssz1Δ* and *zuo1Δ* single deletions strains (Fig. 2B) most probably suggests a ribosome recycling defect. First, it has a similar phenotype to deletions in specific *TMA* genes whose protein products promote 40S subunit recycling, as we and others have previously reported^14,15,21–25^. Second, uORF1 is highly REI permissive thanks to RPEs that functionally interact with the eIF3a/TIF32 subunit of eIF3^20,21,37,38^ to prevent full ribosome recycling. As such, mutations impairing ribosome recycling should show no phenotype with the uORF1 only construct, as was observed (Fig. 2A). Third, the effect of dysregulated -1 or +1 frameshifting can be ruled out, because there are several stop codons in all frames between the main *GCN4* translon and uORF4. Finally, SC-RT can also be discounted because heightened SC-RT decreases and not increases the efficiency of REI past uORF4^19^.

Somewhat unexpectedly, combining *ssz1Δ* and *zuo1Δ* did not further increase the uORF4 reporter activity but virtually nullified the uORF4-specific effect (Fig. 2B). This result indicates that these two factors do not act in a co-operative manner during recycling, even though they can form a heterodimer. The simplest explanation is that each protein independently contributes to controlling a different recycling step. However, to complete the recycling process, the clearance of both independent steps is required, regardless of the order in which it occurs. For example, binding of Ssz1p and Zuo1p to the post-termination ribosome could be mutually exclusive and, therefore, only after the Ssz1p task is completed and Ssz1p leaves, Zuo1p can bind to perform its own task, and *vice versa*. It does not matter which one is done first, both must be performed to avoid blocking the whole process, but they cannot be performed simultaneously. Deleting both factors may modestly reduce the overall recycling rate (neither of them is essential), but at the same time it will remove processivity blocks.

Rai1p has been implicated in mRNA degradation^69^, transcription termination^70–72^, mRNA surveillance and decapping^73,74^, and rRNA processing^40,75^. However, these functions are predominantly bound to the nucleus, often in cooperation with its nuclear binding partner Rat1p. Interestingly, the *rat1* loss-of-function mutation prompts Rai1p to nearly completely re-localize to the cytoplasm^75^, where its roles have not been explored so far. Here, deletion of *rai1* manifested ∼50% decrease in the uORF1-only reporter activity (Fig. 2C), suggesting that one of its cytoplasmic roles could be in promoting REI. The extent of this effect is remarkably similar to those displayed by mutations of two subunits of the eIF3 complex. The *tif35-KLF* mutation of eIF3g reduced the uORF1-only reporter activity down to 52%^76^, and the selected eIF3a mutations reduced it to ∼30-50 %^77^. It is important to emphasize that although it is possible that *rai1Δ* could affect the stability of our reporter mRNAs, as it was suggested to promote mRNA degradation^69^, this potential issue is eliminated by our normalization settings (see Methods). Rai1p and Ssz1p were shown to interact^51^. Interestingly, while combining *ssz1Δ* with *rai1Δ* had no impact on the significantly reduced ORF1 reporter activity observed in the single *rai1Δ* deletion, it produced a compensatory phenotype at uORF4 - increased uORF4 activity of *ssz1Δ* and decreased activity of *rai1Δ* met in the middle and reached nearly wild type values (Fig. 2C and D). This is consistent with the “dominant” REI permissive phenotype of uORF1, where only a defect in REI can manifest itself, but not in recycling, as explained above. However, at uORF4, a defect in REI that further reduces the residual REI capacity of uORF4 may be compensated for by a defect in 40S subunit recycling.

Most importantly, using a simplified uORF1 + uORF4 reporter system and growth assays, we demonstrated that absence of each of these factors diminished the natural inducibility of *GCN4* translation under starvation conditions (by at least 50%) (Fig. 3A), which resulted in a dramatic slow growth phenotype (Fig. 3C). Although the precise molecular mechanism of their action is unclear, we posit that Raip1 contributes to the overall control by promoting efficient REI after uORF1, while Ssz1p and Zuo1p independently ensure complete ribosome recycling after uORF4 translation (Fig. 7). Both of these events are critical for the entire system to operate properly. Interestingly, Ssz1p/Zuo1p have already been implicated in regulation of various stress responses through differing mechanisms, including mito-protein induced stress and heat stress^78^, TOR signaling perturbation^79,80^, arsenite stress^81^, cell wall integrity pathway^80^ or pleiotropic drug resistance^82,83^.

**Figure 7.**
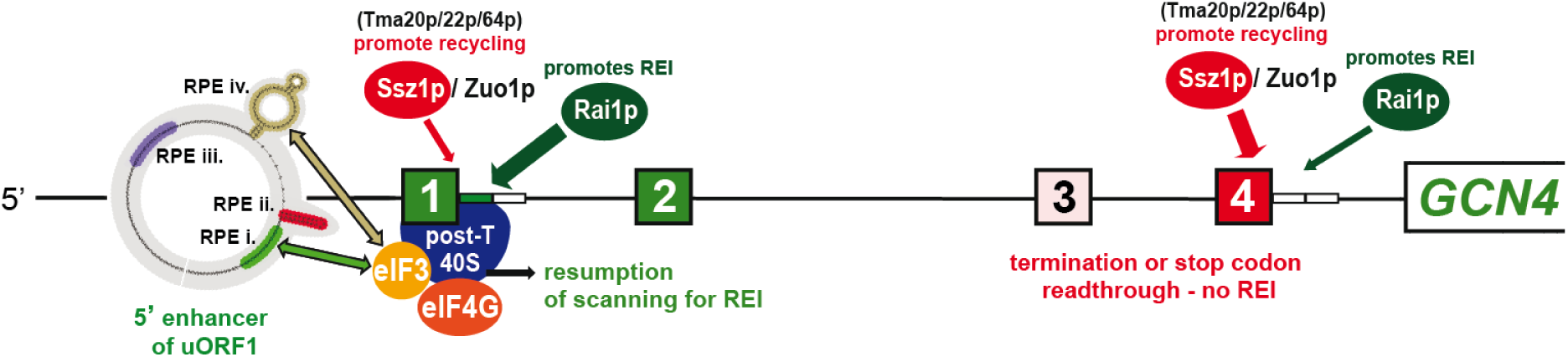
The proposed mechanism of action of Rai1p, Zuo1p, and Ssz1p in *GCN4* translational control. Raip1 contributes to overall control by promoting efficient REI after uORF1, alongside eIF3 and eIF4G, while Ssz1p and Zuo1p independently ensure complete ribosome recycling after uORF4 translation, alongside Tma20p/22p/64p^14,15,21–25^.

We found that Rai1p-FLAG mostly associated with free 40S subunits, and less with 80S monosomes and translating polysomes (Fig. 5A), which is partially at odds with published results^75^, suggesting that that Rai1p binds predominantly to pre-60S particles during 60S biogenesis. However, the authors used cycloheximide to freeze translating ribosomes and admitted that binding to other ribosomal species might occur, but it is below the detection limit of their setup^75^. Our approach was based on crosslinking cells with 1% formaldehyde, which allowed us to detect more transient interactions. Based on our results, we therefore propose that binding of Rai1p to pre-60S does not preclude Rai1p from remaining bound or re-binding to mature ribosomes and thus contributing to translational control.

Our findings combined with the aforementioned fact that Rai1p almost completely re-localizes to the cytoplasm in the *rat1* mutant cells may have interesting implications. It was proposed that Rat1p actively keeps Rai1p tethered to the nucleus to aid in biogenesis. When inactive, Rai1p may migrate to the cytoplasm while bound to immature 60S subunits^75^. Upon amino acid starvation, rRNA transcription is reportedly depleted by ∼80% to downregulate ribosome biogenesis and conserve metabolic energy^84^. This rapid depletion could reduce Rat1p activity and thus allow partial translocation of Rai1p to the cytoplasm to contribute to *GCN4* translational control under stress.

Our stringent co-IP analyses using FLAG-tagged Rai1p as bait followed by LC/MS analysis identified Dbp5p as a major hit for Rai1p, in addition to several ribosomal proteins (Fig. 6A - B). Dbp5p has been proposed to be involved in many facets of RNA surveillance, translation termination and SC-RT^61,85^. Interestingly, among other hits, several enzymes of amino acid biosynthetic pathways were also recorded. Virtually the same applies to Ssz1p-FLAG LC/MS, in addition to the expected enrichment of ribosomal proteins, ribosome- and ribosome biogenesis-associated factors. Both tagged proteins pulled down complexes enriched for Snz1p, which is a *GCN4* transcriptional target gene^62^. The physiological significance of these findings is unclear and is currently under investigation. While Rai1p-FLAG did not pull down Ssz1p and *vice versa*, we detected a direct interaction between these two proteins *in vitro* (Fig. 6C). This is consistent with the proteome survey report^51^. Given that Rai1p interaction with Zuo1p has also been seen^86^, we propose that Rai1p probably coexists with each of its binding partners on the ribosome, but not simultaneously, where they exert antagonistic activities – stimulating REI *versus* ribosome recycling.

In summary, we have identified three new factors contributing to *GCN4* translational control, as well as other yeast mRNAs containing regulatory uTranslons, in the opposite manner. These findings further expand the arsenalof *cis-* and *trans-*acting factors (summarized in Fig. 7) that have been implicated in this delicate, exemplary and intensively-studied mechanism of delayed reinitiation so far. It will be interesting to examine whether their human homologues contribute to *ATF4* translational control in a similar fashion.

## METHODS AND MATERIALS

### β-galactosidase reporter assays

β-galactosidase activities were assayed in whole cell extracts (WCEs) as described in^25^. Briefly, cells were transformed with plasmids bearing *GCN4-lacZ* fusion reporters (along with a *HIS3* carrying shuttle vector for Fig. 3) and grown in synthetic SD media supplemented with histidine, leucine and methionine (for Fig. 3 only leucine and methionine) to mid-log phase. Cells were centrifuged, washed with deionized water and broken using glass beads in Breaking Buffer (100 mM Tris 8.0, 4% Glycerol, 1 mM β-mercaptoethanol), then clarified by centrifugation (18,400*g*/30 min/4 °C). β-galactosidase activity (measured as nanomoles of o-nitrophenyl-3-D-galactose cleaved per minute per milligram of protein) was normalized to protein concentration of the WCE, which was measured by Bradford assay with BSA as standard.

The normalized β-galactosidase activity for each reporter construct in transformants of deletion strains was determined by dividing the unnormalized β-galactosidase activities by a normalization factor. This factor was calculated separately for each individual experiment by assaying the activity of the “uORF-less” reporter (p227) in WT and deletion strains in four transformants under the same conditions as uORF1-only/uORF4-only reporters. Then, the mean of each deletion strain “uORF-less” activities was divided by the mean of WT “uORF-less” activities, determining the normalization factor.

### Plasmids and yeast strains

All plasmids and primers used are listed in Supplementary Tables S2 and S3. Cloning was carried out in the DH5α *E. coli* strain.

The plasmids were constructed as follows: pKJ64 was constructed by replacing the *PstI-BamHI* fragment of YEplac181 by synthesized DNA fragment 1 (GeneString). pKJ65 was constructed by replacing the *HindIII-EcoRI* fragment of YEplac181 by synthesized DNA fragment 2. pKJ72 and pKJ73 were generated by subcloning the synthesized fragments from pKJ64 and pKJ65 into the YCplac111 backbone, as the multicloning site in YEplac181 and YCplac111 is identical. pKJ80 was constructed in two steps: by first, replacing the *PstI-BamHI* fragment of YEplac181 by a fragment amplified from yeast cDNA using primers KJ122 and KJ123, creating plasmid pKJ76 carrying ZUO1 without FLAG. Then, the fragment from pKJ76 was replaced by synthesized DNA fragment 3, creating pKJ79. pKJ80 was generated by sucloning the *PstI-BamHI* fragment from pKJ79. pKJ66 was constructed by replacing the *HindIII-FseI* fragment of pGL4-CMV by a fragment amplified by PCR from yeast cDNA with primers KJ110 and KJ111. pKJ67 was constructed by replacing the *EcoRI-FseI* fragment of pGL4-CMV by a fragment amplified by PCR from yeast cDNA with primers KJ112 and KJ113. pKJ68 was constructed by replacing the *BamHI-SalI* fragment of pGEX-5-X-3 with a fragment amplified by PCR from yeast cDNA using primers KJ108 and KJ109.

### Yeast strain creation

All yeast strains (incl. their genotypes) and primers used are listed in Supplementary Tables S4 and S3.

Yeast deletion strains lacking candidate translation factor genes were sourced from the Euroscarf yeast deletion collection. Yeast double deletion strains were constructed using deletion cassettes carrying the *hphNT1* and *natMX6* genes, carried by plasmids pZC3 and pZC4. The deletion cassettes for *SSZ1* deletion were amplified using PCR from their corresponding plasmids, using primers KJ114 and KJ115. To create YKJ12 (*rai1Δ ssz1Δ*) and YKJ9 (*ssz1Δ zuo1Δ*), the parental yeast strains EUR3 (*rai1Δ*) and EUR78 (*zuo1Δ*) were then transformed with the deletion cassettes targeting *SSZ1* to confer antibiotic resistance and selected using Nourseothricin (Jena Bioscience), or Hygromycin B (Carl Roth GmbH), respectively, on YPD agar medium. Incorporation of the cassettes was verified by colony PCR^87^ using confirmation primers KJ116 and PB238.

### Yeast growth assays

For yeast spot assays, yeast strains were spotted onto minimal synthetic defined (SD) medium plates in five serial 10-fold dilutions (starting with OD_600_ 0.5) and grown for 48 h at 30 °C.

For yeast doubling times, yeast strains were grown in liquid culture overnight to stationary phase, and diluted to OD_600_ = 0.1 the next morning. Diluted cultures were grown in 24 well plates in 1ml of minimal SD media supplemented with Leucine, Methionine and Uracil (additionally, also with 50 mM 3-AT) at 30 °C with high agitation in the Eon™ Microplate Spectrophotometer (BioTek). OD_600_ was measured every 15 minutes. For doubling time calculation, the formula: Doubling Time = (LN(2)/LN(OD_600_ /OD_600_was used. Each experiment was performed in three biological replicates. Each biological replicate was the average of three technical replicates.

### Polysome profiling and fraction collection

For polysome profiling, yeast cultures were grown to OD_600_ ∼ 0.8 in YPD media at 30 °C 230rpm. Then, they were treated with cycloheximide (Sigma Aldrich; 50 μg/1 ml of culture) for 10 minutes and immediately put on ice for 5 minutes. The pellets were washed with dH_2_O and then resuspended in 1 ml of GA^+^ buffer (20 mM Tris-HCl (pH 7.5), 50 mM KCl, 10 mM MgCl2, 1 mM DTT, 5 mM NaF, 50 μg/ml cycloheximide). After centrifugation, the pellet was again resuspended in 300 μl of GF buffer (20 mM Tris-HCl (pH 7.5), 50 mM KCl, 10 mM MgCl2, 1 mM DTT, 5 mM NaF, 50 μg/ml cycloheximide, C0mplete EDTA-free Protease Inhibitor mix (Roche)) and cells were broken using glass beads. Afterwards, the lysate was clarified at 13,500g/10 minutes/4 °C. 8 AU OD_260_ were loaded on a 5-50% sucrose gradient and separated by ultracentrifugation at 39,000rpm for 2.5 hours using the SW41Ti rotor. The gradients were scanned at A254 nm using the Biocomp Gradient Station™ (Biocomp Instruments).

For fraction collection, yeast strains transformed with FLAG-Ssz1p and FLAG-Rai1p plasmids were grown in minimal SD media supplemented with Histidine, Methionine and Uracil to OD_600_ ∼ 0.9-1.0 at 30 °C 230rpm. Then, cells were put on ice and 1% formaldehyde (Sigma Aldrich) was added as a crosslinking agent and incubated for 60 minutes on ice. Afterwards, 10 ml of 2.5M Glycine/100 ml of culture was added. The pellets were then resuspended in GF buffer, centrifuged, again resuspended in 600μl of GF buffer and broken using glass beads. 20 AU OD_260_ were loaded on a 5-50% sucrose gradient and separated by ultracentrifugation at 39,000rpm for 2.5 hours using the SW41Ti rotor. Fractions were collected using the Biocomp Gradient Station™ (Biocomp Instruments). Each fraction was precipitated with 96% Ethanol (2x volume of the fraction) at -20 °C overnight, then centrifuged at 21,000g/1 hour/4 °C. The resulting pellets were then resuspended in 100 μl of 1x SDS-PAGE Sample Buffer and 30 μl/well was used for western blot analysis. 30 μl of WCE was used as input.

### Western Blot

Samples were resolved through SDS-PAGE followed by western blotting. For SDS-PAGE, Criterion TGX™ Gels were used. After wet transfer, membranes were blocked in 5% milk for 60 minutes and incubated with primary antibodies at 4 °C overnight. The primary antibodies used were anti-FLAG M2 (Sigma Aldrich, cat# F1804, lot# 080M6035; dilution 1:500) and anti-TIF32 (sourced from Lon Phan, NCBI, USA; dilution 1:1000). Membranes were washed 3x5 minutes in TBS-T buffer and then incubated in secondary antibodies at room temperature for 90 minutes. The secondary antibodies used were Amersham ECL™ Anti-Mouse IgG HRP-linked (Cytiva, cat# NA931V, lot# 17853437, dilution 1:2000) and Amersham ECL™ Anti-Rabbit IgG HRP-linked (Cytiva, cat# NA934V, lot# 18227506, dilution 1:2500). Afterwards, the membranes were washed 3x5 minutes in TBS-T and the signal was developed using SuperSignal West Femto Maximum Sensitivity Substrate (Thermo Fisher Scientific) and imaged using G-Box (Syngene).

### Co-Immunoprecipitation

Yeast strain *rai1Δ/ssz1Δ* was transformed with FLAG-Rai1/FLAG-SSZ1 plasmid, then grown along with *rai1Δ/ssz1Δ* (no plasmid) in SD media supplemented with Histidine, Methionine and Uracil (and Leucine) to OD_600_ ∼ 0.8 at 30 °C 230rpm. The pellets were washed with 25 ml of cold FLAG buffer (20 mM TRIS (pH 7.5), 200 mM KCl, 5 mM MgAc), then weighed and resuspended in cold FLAG buffer with protease inhibitors (Aprotinin 1μg/ml, Leupeptin 1μg/ml, Pepstatin 1μg/ml, 1 mM PMSF, C0mplete EDTA-free Protease Inhibitor mix (Roche), 1 mM DTT). The amount of buffer used was 1 ml/g of pellet weight. The cells were broken using glass beads and the lysate was subsequently cleared 2x at 18,400g/10 min/4 °C. For the pulldown, ANTI-FLAG® M2 Affinity Gel (Milipore) beads slurry was washed 3x with cold FLAG buffer (with protease inhibitors). Afterwards, whole cell lysate was added (0.2mg of total protein adjusted to 1ml volume with FLAG buffer) to the beads (2.5/10 μl of beads slurry for Rai1/Ssz1, respectively) and incubated for 2 hours at 4 °C on a rotator. Then, the beads were washed 3x with 1 ml of cold FLAG buffer with 0.2% Triton X-100 (Sigma Aldrich), 1x with cold FLAG buffer without Triton X-100 and 2x with cold 50 mM TRIS (pH 8.0) and finally resuspended in 50μl of 50 mM TRIS (pH 8.0) to be sent for MS analysis. Total protein concentrations were determined using Bradford assay.

### Protein Digestion

Beads were resuspended with 100 ul of 100mM TEAB containing 2% SDC. Cysteins were reduced with 10mM final concentration of TCEP and blocked with 40mM final concentration of chloroacetamide (60°C for 30 min). Samples were cleaved on beads with 1 µg of trypsine at 37°C overnight. After digestion samples were centrifuged and supernatants were collected and acidified with TFA to 1% final concentration. SDC was removed by extraction to ethylacetate^88^. Peptides were desalted using in-house made stage tips packed with C18 disks (Empore) according to ^89^.

### nLC-MS 2 Analysis

Nano Reversed phase column (Ion Opticks Aurora Ultimate XT 25 cm x 75 µm ID, C18 UHPLC column, 1.7 µm particles, 120 Å pore size) was used for LC/MS analysis. Mobile phase buffer A was composed of water and 0.1% formic acid. Mobile phase B was composed of acetonitrile and 0.1% formic acid. Samples were loaded onto the trap column (Acclaim PepMap300, C18, 5 µm, 300 Å Wide Pore, 300 µm x 5 mm, 5 Cartridges) for 4 min at 15 μl/min. Loading buffer was composed of water, 2% acetonitrile and 0.1% trifluoroacetic acid. Peptides were eluted with Mobile Phase B gradient from 4% to 35% B in 60 min. Eluting peptide cations were converted to gas-phase ions by electrospray ionization and analysed on a Thermo Orbitrap Ascend (Q-OT- qIT, Thermo). Survey scans of peptide precursors from 350 to 1400 m/z were performed at 120K resolution (at 200 m/z) with a 5 × 10^5^ ion count target. Tandem MS was performed by isolation at 1,5 Th with the quadrupole, HCD fragmentation with normalized collision energy of 30, and rapid scan MS analysis in the ion trap. The MS/MS ion count target was set to 10^4^ and the max injection time was 35 ms. Only those precursors with charge state 2–6 were sampled for MS/MS. The dynamic exclusion duration was set to 45 s with a 10 ppm tolerance around the selected precursor and its isotopes. Monoisotopic precursor selection was turned on. The instrument was run in top speed mode with 2 s cycles^90^.

### Protein expression and purification

The E.coli strain BL21 Star (DE3) (Invitrogen) was used for expression and purification of GST-tagged Ssz1 protein. Expression of GST-Ssz1 was induced by 10 mM IPTG (Serva) overnight at 16 °C 230rpm in LB media. Pellet was washed in cold dH_2_O and resuspended in 5 ml of cold PBS (pH 7.5, 137 mM NaCl, 2.7 mM KCl, 10 mM Na2HPO4, 1.8 mM KH2PO4). Cells were broken using glass beads and 1.5 % Triton X-100 (Sigma Aldrich) was added. The mixture was incubated on a rotator for 30 minutes at 4 °C and then centrifuged at 21,000g/10 min/4°C. Gluthathione Separose beads (50% slurry; GE Healthcare) were washed 3x with PBS and 400μl of the slurry was added into the clarified cell lysate. The lysate and beads were then incubated for 30 minutes at RT. Afterwards, the beads were washed 3x with cold PBS, aliquoted and frozen at -80 °C for use in the GST pulldown.

### In vitro GST pulldown

^35^S-labelled proteins were produced using the TnT® Quick Coupled Transcription/Translation System (Promega) according to vendor instructions. GST-Ssz1 on Separose beads was incubated with ^35^S-Rai1 and ^35^S-Zuo1 for 2 hours at 4 °C in 300μl of Buffer B (20 mM HEPES (pH 7.5), 75 mM KCl, 0.1 mM EDTA, 2.5 mM MgCl2, 0.05% Igepal CA-630, 1mM DTT, 1% fat-free milk) and then washed 3x with PBS. The proteins were separated by SDS-PAGE, gels were first stained with Simply Blue™ SafeStain (Invitrogen) and then subjected to autoradiography.

### Statistical analysis

For all reporter assays, all data were tested for normality by the Shapiro–Wilk test. After confirming normality, the parametric unpaired, two-tailed Welch’s *t* test was used; **** indicates *p* < 0.0001; \*\*\**p* < 0.001; \*\**p* < 0.01; \**p* < 0.05; ns non-significant. All individual data points are shown as part of bar plots. The SD was calculated from all biological replicates for each construct in each strain. Statistical analysis and visualization were performed in GraphPad Prism, version 9.4.1 (GraphPad Software). All replicates shown in the reporter assays graphs were independent transformants—biological replicates.

All MS data were analyzed and quantified with the MaxQuant software (version 2.3.1.0)^91^. The false discovery rate (FDR) was set to 1% for both proteins and peptides and we specified a minimum peptide length of seven amino acids. The Andromeda search engine was used for the MS/MS spectra search against the Saccharomyces cerevisiae database (downloaded from Uniprot.org in February 2024, containing 6 060 entries). Enzyme specificity was set as C-terminal to Arg and Lys, also allowing cleavage at proline bonds and a maximum of two missed cleavages. Carbmaidomethylation of cysteine was selected as fixed modification and N- terminal protein acetylation and methionine oxidation as variable modifications. Data analysis was performed using Perseus 1.6.15.0 software.^92^

For volcano plot creation, the VolcaNoseR tool was used^93^. Log_2_FC threshold was set as -1 and +1 (corresponding to 2-fold change) and threshold –Log p value was set as 2 (corresponding to p = 0.01).

## ACKNOWLEDGEMENTS

We thank members of the Laboratory of Regulation of Gene Expression for their fruitful discussion. In particular, we thank Stanislava Gunisova for help with the set-up of the initial screening strategy. LC-MS analyses were performed in OMICS Mass Spectrometry Core Facility at Biocev research center, Faculty of Science, Charles University. This work was supported by the Czech Science Foundation grant 24-10013S, the Praemium Academiae grant provided by the Czech Academy of Sciences, and CZ.02.01.01/00/22_008/0004575 RNA for therapy by ERDF and MEYS (all to LSV).

## DATA AVAILABILITY STATEMENT

All data from this study are available within this paper and its supplementary information. All plasmids and strains used in this study are available upon request from the authors. Source data for graphs presented in the main figures can be found in the Supplementary Data 1 file.

## AUTHOR CONTRIBUTION

K.J. and L.S.V. conceived and designed the project.

K.J. carried out majority of experiments and performed the data analysis; she was assisted by A.S.

K.J. and L.S.V. interpreted the results.

K.J. and L.S.V. wrote the paper.

---

**Figure S1.**
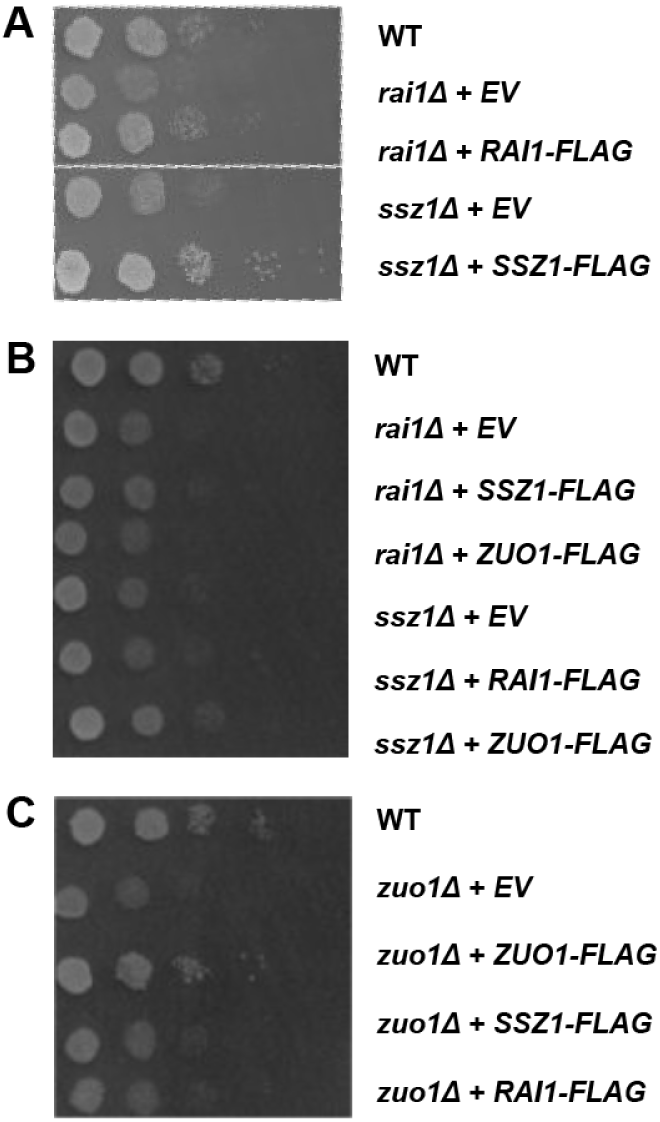
Growth complementation spot assays. **A - C.** WT, *rai1Δ*, *ssz1Δ* and *zuo1Δ* strains were transformed with either the EV (YEplac181) or YEplac181 plasmids carrying either *SSZ1-FLAG, ZUO1-FLAG or* or *RAI1-FLAG* fusions. The cultures were first diluted to an OD_600_ = 0.5, then serially diluted (10x) and spotted onto SD plates. The plates were incubated at 30 °C for 48 hours.

## REFERENCES

1. Hinnebusch, A. G. Structural Insights into the Mechanism of Scanning and Start Codon Recognition in Eukaryotic Translation Initiation. Trends Biochem. Sci. 42, 589–611 (2017).

2. Brito Querido, J., Díaz-López, I. & Ramakrishnan, V. The molecular basis of translation initiation and its regulation in eukaryotes. Nat. Rev. Mol. Cell Biol. 25, 168–186 (2024).

3. Dever, T. E., Ivanov, I. P. & Hinnebusch, A. G. Translational regulation by uORFs and start codon selection stringency. Genes Dev. 37, 474–489 (2023).

4. Świrski, M. I., et al. Translon: a single term for translated regions. Nat. Methods 22, 2002–2006 (2025).

5. Gunišová, S., Hronová, V., Mohammad, M. P., Hinnebusch, A. G. & Valášek, L. S. Please do not recycle! Translation reinitiation in microbes and higher eukaryotes. FEMS Microbiol. Rev. 42, 165–192 (2017).

6. Calvo, S. E., Pagliarini, D. J. & Mootha, V. K. Upstream open reading frames cause widespread reduction of protein expression and are polymorphic among humans. Proc. Natl. Acad. Sci. U. S. A. 106, 7507–7512 (2009).

7. Lawless, C. et al. Upstream sequence elements direct post-transcriptional regulation of gene expression under stress conditions in yeast. BMC Genomics 10, 7 (2009).

8. Skabkin, M. A. et al. Activities of Ligatin and MCT-1/DENR in eukaryotic translation initiation and ribosomal recycling. Genes Dev. 24, 1787–1801 (2010).

9. Ahmed, Y. L. et al. DENR–MCTS1 heterodimerization and tRNA recruitment are required for translation reinitiation. PLoS Biol. 16, e2005160 (2018).

10. Hellen, C. U. T. Translation Termination and Ribosome Recycling in Eukaryotes. Cold Spring Harb. Perspect. Biol. 10, a032656 (2018).

11. Luukkonen, B. G., Tan, W. & Schwartz, S. Efficiency of reinitiation of translation on human immunodeficiency virus type 1 mRNAs is determined by the length of the upstream open reading frame and by intercistronic distance. J. Virol. 69, 4086–4094 (1995).

12. Kozak, M. Constraints on reinitiation of translation in mammals. Nucleic Acids Res. 29, 5226–5232 (2001).

13. Rajkowitsch, L., Vilela, C., Berthelot, K., Ramirez, C. V. & McCarthy, J. E. G. Reinitiation and recycling are distinct processes occurring downstream of translation termination in yeast. J. Mol. Biol. 335, 71–85 (2004).

14. Mohammad, M. P., Munzarová Pondelícková, V., Zeman, J., Gunišová, S. & Valášek, L. S. In vivo evidence that eIF3 stays bound to ribosomes elongating and terminating on short upstream ORFs to promote reinitiation. Nucleic Acids Res. 45, 2658–2674 (2017).

15. Mohammad, M. P., Smirnova, A., Gunišová, S. & Valášek, L. S. eIF4G is retained on ribosomes elongating and terminating on short upstream ORFs to control reinitiation in yeast. Nucleic Acids Res. 49, 8743–8756 (2021).

16. Bohlen, J., Fenzl, K., Kramer, G., Bukau, B. & Teleman, A. A. Selective 40S Footprinting Reveals Cap-Tethered Ribosome Scanning in Human Cells. Mol. Cell 79, 561–574.e5 (2020).

17. Wagner, S. et al. Selective Translation Complex Profiling Reveals Staged Initiation and Co-translational Assembly of Initiation Factor Complexes. Mol. Cell 79, 546–560.e7 (2020).

18. Grant, C. M., Miller, P. F. & Hinnebusch, A. G. Requirements for intercistronic distance and level of eukaryotic initiation factor 2 activity in reinitiation on GCN4 mRNA vary with the downstream cistron. Mol. Cell. Biol. 14, 2616–2628 (1994).

19. Gunišová, S., Beznosková, P., Mohammad, M. P., Vlčková, V. & Valášek, L. S. In-depth analysis of cis-determinants that either promote or inhibit reinitiation on GCN4 mRNA after translation of its four short uORFs. RNA N. Y. N 22, 542–558 (2016).

20. Munzarová, V. et al. Translation reinitiation relies on the interaction between eIF3a/TIF32 and progressively folded cis-acting mRNA elements preceding short uORFs. PLoS Genet. 7, e1002137 (2011).

21. Szamecz, B. et al. eIF3a cooperates with sequences 5’ of uORF1 to promote resumption of scanning by post-termination ribosomes for reinitiation on GCN4 mRNA. Genes Dev. 22, 2414–2425 (2008).

22. Young, D. J. et al. Tma64/eIF2D, Tma20/MCT-1, and Tma22/DENR Recycle Post-termination 40S Subunits In Vivo. Mol. Cell 71, 761–774.e5 (2018).

23. Bohlen, J. et al. DENR promotes translation reinitiation via ribosome recycling to drive expression of oncogenes including ATF4. Nat. Commun. 11, 4676 (2020).

24. Young, D. J., Meydan, S. & Guydosh, N. R. 40S ribosome profiling reveals distinct roles for Tma20/Tma22 (MCT-1/DENR) and Tma64 (eIF2D) in 40S subunit recycling. Nat. Commun. 12, 2976 (2021).

25. Jendruchová, K. et al. Differential effects of 40S ribosome recycling factors on reinitiation at regulatory uORFs in GCN4 mRNA are not dictated by their roles in bulk 40S recycling. *Commun*. Biol. 7, 1083 (2024).

26. Hinnebusch, A. G. Translational regulation of GCN4 and the general amino acid control of yeast. Annu. Rev. Microbiol. 59, 407–450 (2005).

27. Hinnebusch, A. G. Mechanisms of gene regulation in the general control of amino acid biosynthesis in Saccharomyces cerevisiae. Microbiol. Rev. 52, 248–273 (1988).

28. Mueller, P. P. & Hinnebusch, A. G. Multiple upstream AUG codons mediate translational control of GCN4. Cell 45, 201–207 (1986).

29. Dever, T. E. et al. Phosphorylation of initiation factor 2 alpha by protein kinase GCN2 mediates gene-specific translational control of GCN4 in yeast. Cell 68, 585–596 (1992).

30. Pakos-Zebrucka, K. et al. The integrated stress response. EMBO Rep. 17, 1374–1395 (2016).

31. Smirnova, A. M. et al. Stem-loop-induced ribosome queuing in the uORF2/ATF4 overlap fine-tunes stress-induced human ATF4 translational control. Cell Rep. 43, 113976 (2024).

32. Andreev, D. E. et al. Translation of 5’ leaders is pervasive in genes resistant to eIF2 repression. eLife 4, e03971 (2015).

33. Sidrauski, C., McGeachy, A. M., Ingolia, N. T. & Walter, P. The small molecule ISRIB reverses the effects of eIF2α phosphorylation on translation and stress granule assembly. eLife 4, e05033 (2015).

34. Starck, S. R. et al. Translation from the 5′ untranslated region shapes the integrated stress response. Science 351, aad3867 (2016).

35. Chen, C.-W. et al. Plasticity of the mammalian integrated stress response. Nature 641, 1319–1328 (2025).

36. Guan, B.-J. et al. A Unique ISR Program Determines Cellular Responses to Chronic Stress. Mol. Cell 68, 885–900.e6 (2017).

37. Gunišová, S. & Valášek, L. S. Fail-safe mechanism of GCN4 translational control--uORF2 promotes reinitiation by analogous mechanism to uORF1 and thus secures its key role in GCN4 expression. Nucleic Acids Res. 42, 5880–5893 (2014).

38. Grant, C. M., Miller, P. F. & Hinnebusch, A. G. Sequences 5’ of the first upstream open reading frame in GCN4 mRNA are required for efficient translational reinitiation. Nucleic Acids Res. 23, 3980–3988 (1995).

39. Grant, C. M. & Hinnebusch, A. G. Effect of sequence context at stop codons on efficiency of reinitiation in GCN4 translational control. Mol. Cell. Biol. 14, 606–618 (1994).

40. Fang, F., Phillips, S. & Butler, J. S. Rat1p and Rai1p function with the nuclear exosome in the processing and degradation of rRNA precursors. RNA 11, 1571–1578 (2005).

41. Doamekpor, S. K. et al. DXO/Rai1 enzymes remove 5′-end FAD and dephospho-CoA caps on RNAs. Nucleic Acids Res. 48, 6136–6148 (2020).

42. Xue, Y. et al. Saccharomyces cerevisiae RAI1 (YGL246c) Is Homologous to Human DOM3Z and Encodes a Protein That Binds the Nuclear Exoribonuclease Rat1p. Mol. Cell. Biol. 20, 4006 (2000).

43. Kišonaitė, M. et al. Structural inventory of cotranslational protein folding by the eukaryotic RAC complex. Nat. Struct. Mol. Biol. 30, 670–677 (2023).

44. Gautschi, M. et al. RAC, a stable ribosome-associated complex in yeast formed by the DnaK-DnaJ homologs Ssz1p and zuotin. Proc. Natl. Acad. Sci. U. S. A. 98, 3762–3767 (2001).

45. Lee, K., Sharma, R., Shrestha, O. K., Bingman, C. A. & Craig, E. A. Dual interaction of the Hsp70 J protein co-chaperone Zuotin with the 40S and 60S subunits of the ribosome. Nat. Struct. Mol. Biol. 23, 1003–1010 (2016).

46. Leidig, C. et al. Structural characterization of a eukaryotic chaperone—the ribosome-associated complex. Nat. Struct. Mol. Biol. 20, 23–28 (2013).

47. Rakwalska, M. & Rospert, S. The Ribosome-Bound Chaperones RAC and Ssb1/2p Are Required for Accurate Translation in Saccharomyces cerevisiae. Mol. Cell. Biol. 24, 9186–9197 (2004).

48. Grant, C. M., Miller, P. F. & Hinnebusch, A. G. Sequences 5’ of the first upstream open reading frame in GCN4 mRNA are required for efficient translational reinitiation. Nucleic Acids Res. 23, 3980–3988 (1995).

49. Brachmann, C. B. et al. Designer deletion strains derived from Saccharomyces cerevisiae S288C: a useful set of strains and plasmids for PCR-mediated gene disruption and other applications. Yeast Chichester Engl. 14, 115–132 (1998).

50. Fleischer, T. C., Weaver, C. M., McAfee, K. J., Jennings, J. L. & Link, A. J. Systematic identification and functional screens of uncharacterized proteins associated with eukaryotic ribosomal complexes. Genes Dev. 20, 1294–1307 (2006).

51. Gavin, A.-C. et al. Proteome survey reveals modularity of the yeast cell machinery. Nature 440, 631–636 (2006).

52. Klopotowski, T. & Wiater, A. Synergism of aminotriazole and phosphate on the inhibition of yeast imidazole glycerol phosphate dehydratase. Arch. Biochem. Biophys. 112, 562–566 (1965).

53. Alifano, P. et al. Histidine biosynthetic pathway and genes: structure, regulation, and evolution. Microbiol. Rev. 60, 44–69 (1996).

54. Wek, S. A., Zhu, S. & Wek, R. C. The histidyl-tRNA synthetase-related sequence in the eIF-2 alpha protein kinase GCN2 interacts with tRNA and is required for activation in response to starvation for different amino acids. Mol. Cell. Biol. 15, 4497–4506 (1995).

55. Chen, Y., Tsai, B., Li, N. & Gao, N. Structural remodeling of ribosome associated Hsp40-Hsp70 chaperones during co-translational folding. Nat. Commun. 13, 3410 (2022).

56. Valásek, L., Trachsel, H., Hasek, J. & Ruis, H. Rpg1, the Saccharomyces cerevisiae Homologue of the Largest Subunit of Mammalian Translation Initiation Factor 3, Is Required for Translational Activity *. J. Biol. Chem. 273, 21253–21260 (1998).

57. Valásek, L. et al. The yeast eIF3 subunits TIF32/a, NIP1/c, and eIF5 make critical connections with the 40S ribosome in vivo. Genes Dev. 17, 786–799 (2003).

58. Beznosková, P. et al. Translation Initiation Factors eIF3 and HCR1 Control Translation Termination and Stop Codon Read-Through in Yeast Cells. PLoS Genet. 9, e1003962 (2013).

59. Beznosková, P., Wagner, S., Jansen, M. E., von der Haar, T. & Valášek, L. S. Translation initiation factor eIF3 promotes programmed stop codon readthrough. Nucleic Acids Res. 43, 5099–5111 (2015).

60. Mellacheruvu, D. et al. The CRAPome: a contaminant repository for affinity purification-mass spectrometry data. Nat. Methods 10, 730–736 (2013).

61. Querl, L. & Krebber, H. The DEAD-box RNA helicase Dbp5 is a key protein that couples multiple steps in gene expression. Biol. Chem. 404, 845–850 (2023).

62. Qiu, H. et al. An array of coactivators is required for optimal recruitment of TATA binding protein and RNA polymerase II by promoter-bound Gcn4p. Mol. Cell. Biol. 24, 4104–4117 (2004).

63. Gautschi, M. et al. RAC, a stable ribosome-associated complex in yeast formed by the DnaK-DnaJ homologs Ssz1p and zuotin. Proc. Natl. Acad. Sci. U. S. A. 98, 3762–3767 (2001).

64. Huang, P., Gautschi, M., Walter, W., Rospert, S. & Craig, E. A. The Hsp70 Ssz1 modulates the function of the ribosome-associated J-protein Zuo1. Nat. Struct. Mol. Biol. 12, 497–504 (2005).

65. Zhang, Y. et al. The ribosome-associated complex RAC serves in a relay that directs nascent chains to Ssb. Nat. Commun. 11, 1504 (2020).

66. Kišonaitė, M. et al. Structural inventory of cotranslational protein folding by the eukaryotic RAC complex. Nat. Struct. Mol. Biol. 30, 670–677 (2023).

67. Muldoon-Jacobs, K. L. & Dinman, J. D. Specific effects of ribosome-tethered molecular chaperones on programmed -1 ribosomal frameshifting. Eukaryot. Cell 5, 762–770 (2006).

68. Jaiswal, H. et al. The Chaperone Network Connected to Human Ribosome-Associated Complex. Mol. Cell. Biol. 31, 1160–1173 (2011).

69. Das, B., Butler, J. S. & Sherman, F. Degradation of Normal mRNA in the Nucleus of Saccharomyces cerevisiae. Mol. Cell. Biol. 23, 5502–5515 (2003).

70. Kim, M. et al. The yeast Rat1 exonuclease promotes transcription termination by RNA polymerase II. Nature 432, 517–522 (2004).

71. El Hage, A., Koper, M., Kufel, J. & Tollervey, D. Efficient termination of transcription by RNA polymerase I requires the 5’ exonuclease Rat1 in yeast. Genes Dev. 22, 1069–1081 (2008).

72. Park, J., Kang, M. & Kim, M. Unraveling the mechanistic features of RNA polymerase II termination by the 5’-3’ exoribonuclease Rat1. Nucleic Acids Res. 43, 2625–2637 (2015).

73. Wang, V. Y.-F., Jiao, X., Kiledjian, M. & Tong, L. Structural and biochemical studies of the distinct activity profiles of Rai1 enzymes. Nucleic Acids Res. 43, 6596–6606 (2015).

74. Zhang, Y. et al. Extensive 5’-surveillance guards against non-canonical NAD-caps of nuclear mRNAs in yeast. Nat. Commun. 11, 5508 (2020).

75. Sydorskyy, Y. et al. Intersection of the Kap123p-Mediated Nuclear Import and Ribosome Export Pathways. Mol. Cell. Biol. 23, 2042–2054 (2003).

76. Cuchalová, L. et al. The RNA Recognition Motif of Eukaryotic Translation Initiation Factor 3g (eIF3g) Is Required for Resumption of Scanning of Posttermination Ribosomes for Reinitiation on GCN4 and Together with eIF3i Stimulates Linear Scanning. Mol. Cell. Biol. 30, 4671–4686 (2010).

77. Munzarová, V. et al. Translation reinitiation relies on the interaction between eIF3a/TIF32 and progressively folded cis-acting mRNA elements preceding short uORFs. PLoS Genet. 7, e1002137 (2011).

78. Qian, J. et al. The ribosome-associated complex regulates cytosolic translation upon mitoprotein-induced stress. FEBS J. 10.1111/febs.70356 (2025) doi:10.1111/febs.70356.

79. Black, A. et al. The ribosome-associated chaperone Zuo1 controls translation upon TORC1 inhibition. EMBO J. 42, e113240 (2023).

80. Gillies, A. T., Taylor, R. & Gestwicki, J. E. Synthetic lethal interactions in yeast reveal functional roles of J protein co-chaperones. Mol. Biosyst. 8, 2901–2908 (2012).

81. Rodrigues, J. I., Lorentzon, E., Hua, S., Boucher, A. & Tamás, M. J. Yeast chaperones and ubiquitin ligases contribute to proteostasis during arsenite stress by preventing or clearing protein aggregates. FEBS Lett. 597, 1733–1747 (2023).

82. Eisenman, H. C. & Craig, E. A. Activation of pleiotropic drug resistance by the J-protein and Hsp70-related proteins, Zuo1 and Ssz1. Mol. Microbiol. 53, 335–344 (2004).

83. Prunuske, A. J., Waltner, J. K., Kuhn, P., Gu, B. & Craig, E. A. Role for the molecular chaperones Zuo1 and Ssz1 in quorum sensing via activation of the transcription factor Pdr1. Proc. Natl. Acad. Sci. U. S. A. 109, 472–477 (2012).

84. Conesa, C. et al. Modulation of yeast genome expression in response to defective RNA polymerase III-dependent transcription. Mol. Cell. Biol. 25, 8631–8642 (2005).

85. Rajan, A. A. N. & Montpetit, B. Emerging molecular functions and novel roles for the DEAD-box protein Dbp5/DDX19 in gene expression. Cell. Mol. Life Sci. CMLS 78, 2019 (2020).

86. Michaelis, A. C. et al. The social and structural architecture of the yeast protein interactome. Nature 624, 192–200 (2023).

87. Lõoke, M., Kristjuhan, K. & Kristjuhan, A. EXTRACTION OF GENOMIC DNA FROM YEASTS FOR PCR-BASED APPLICATIONS. BioTechniques 50, 325–328 (2011).

88. Masuda, T., Tomita, M. & Ishihama, Y. Phase transfer surfactant-aided trypsin digestion for membrane proteome analysis. J. Proteome Res. 7, 731–740 (2008).

89. Rappsilber, J., Mann, M. & Ishihama, Y. Protocol for micro-purification, enrichment, pre-fractionation and storage of peptides for proteomics using StageTips. Nat. Protoc. 2, 1896–1906 (2007).

90. Hebert, A. S. et al. The one hour yeast proteome. Mol. Cell. Proteomics MCP 13, 339–347 (2014).

91. Cox, J. et al. Accurate proteome-wide label-free quantification by delayed normalization and maximal peptide ratio extraction, termed MaxLFQ. Mol. Cell. Proteomics MCP 13, 2513–2526 (2014).

92. Tyanova, S. et al. The Perseus computational platform for comprehensive analysis of (prote)omics data. Nat. Methods 13, 731–740 (2016).

93. Goedhart, J. & Luijsterburg, M. S. VolcaNoseR is a web app for creating, exploring, labeling and sharing volcano plots. Sci. Rep. 10, 20560 (2020).

